# Pesticide exposure affects flight dynamics and reduces flight endurance in bumblebees

**DOI:** 10.1101/449280

**Authors:** Daniel Kenna, Hazel Cooley, Ilaria Pretelli, Ana Ramos Rodrigues, Steve D. Gill, Richard J. Gill

**Affiliations:** Department of Life Sciences, Imperial College London, Silwood Park, Ascot, Berkshire, SL5 7PY, UK; Dipartimento di Biologia, Università di Padova, 35121 Padova, Italy; Department of Human Behaviour, Ecology, and Culture, Max Planck Institute for Evolutionary Anthropology, 04103 Leipzig, Germany

**Keywords:** *Bombus terrestris audax*, flight mill, foraging, Imidacloprid, neonicotinoid, velocity

## Abstract

The emergence of agricultural land use change creates a number of challenges that insect pollinators, such as eusocial bees, must overcome. Resultant fragmentation and loss of suitable foraging habitats, combined with pesticide exposure, may increase demands on foraging, specifically the ability to reach resources under such stress. Understanding the effect that pesticides have on flight performance is therefore vital if we are to assess colony success in these changing landscapes. Neonicotinoids are one of the most widely used classes of pesticide across the globe, and exposure to bees has been associated with reduced foraging efficiency and homing ability. One explanation for these effects could be that elements of flight are being affected, but apart from a couple of studies on the honeybee, this has scarcely been tested. Here we used flight mills to investigate how exposure to a field realistic (10ppb) acute dose of imidacloprid affected flight performance of a wild insect pollinator - the bumblebee, *Bombus terrestris audax*. Intriguingly, intial observations showed exposed workers flew at a significantly higher velocity over the first ¾ km of flight. This apparent hyperactivity, however, may have a cost as exposed workers showed reduced flight distance and duration to around a third of what control workers were capable of achieving. Given that bumblebees are central place foragers, impairment to flight endurance could translate to a decline in potential forage area, decreasing the abundance, diversity and nutritional quality of available food, whilst potentially diminishing pollination service capabilities.

**Summary Statement:** Acute neonicotinoid exposure impaired flight endurance and affected velocity of *Bombus terrestris* workers, which may dramatically reduce colony foraging potential and pollination provision in pesticide applied landscapes.

## Introduction

The extent to which insects move across landscapes has significant implications for human welfare. Highly mobile species can potentially lead to detrimental insect pest outbreaks (Mazzi and Dorn, 2012; Sharov and Liebhold, 1998), invasions (Myers et al., 2000; Renault et al., 2018) or the spread of vector borne diseases (Dujardin et al., 2008; Estrada-Peña et al., 2014; Githeko et al., 2000; Rogers and Packer, 1993). Yet insect movement can also underpin beneficial ecosystem service provision. For example, the majority of angiosperms, including around ¾ of our crop species, are to some degree reliant upon the extensive movement of foraging insect pollinators (Gill et al., 2016; Kleijn et al., 2015; Klein et al., 2007; Ollerton et al., 2011). It is therefore important we understand which, and to what extent, stressors can impact on insect pollinator flight performance if we are to mitigate threats to a global pollination service valued at >€150bn annually (Benaets et al., 2017; Fischer et al., 2014; Gallai et al., 2009; Gill and Raine, 2014; Stanley et al., 2015; Wolf et al., 2014).

The emergence of intensive agriculture can cause loss and fragmentation of suitable foraging habitats, leading to resources becoming increasingly sparse and isolated within an insect’s foraging range (Didham et al., 1996; Hadley and Betts, 2012; Steffan-Dewenter and Tscharntke, 1999; Tscharntke and Brandl, 2004; Zurbuchen et al., 2010). This may pose a considerable challenge for eusocial bees, which are characterised as central place foragers by having a fixed nest site. Workers must undertake return foraging trips from this set nest location, and consequently any habitat discontinuity may require workers to fly longer distances to find and bring back resources, such as pollen and nectar (Goulson et al., 2008; Jha and Kremen, 2013; Pelletier and McNeil, 2003; Schmid-Hempel and Schmid-Hempel, 1998). Hence any stressor lowering individual worker flight ability could translate to negative colony level impacts (Gill et al., 2012), with implications for the crucial ecosystem services they provide (Delaplane and Mayer, 2000; Garibaldi et al., 2013; Greenleaf and Kremen, 2006; Potts et al., 2010; Winfree et al., 2008).

Insecticides are commonly applied in agricultural landscapes as a pest management strategy (Fernandez-Cornejo and Vialou, 2014; Ramankutty et al., 2018), with neonicotinoids being one of the most widely used classes worldwide (Simon-Delso et al., 2015). However, neonicotinoids have been implicated as a threat to eusocial bees (Gill et al., 2012; Goulson, 2013; Lundin et al., 2015; Tsvetkov et al., 2017; Whitehorn et al., 2012; Woodcock et al., 2017). Foraging eusocial bees are known to be frequently exposed to neonicotinoids in treated landscapes (Botías et al., 2015; Botías et al., 2017; David et al., 2016; Mitchell et al., 2017; Rolke et al., 2016), and controlled exposure experiments have demonstrated both impaired homing ability (Fischer et al., 2014; Stanley et al., 2016) and foraging efficiency of workers, including longer foraging trips and reduced rate of pollen collection (Feltham et al., 2014; Gill and Raine, 2014; Stanley and Raine, 2016). A possible explanation for these reported impairments is that certain aspects of foraging flight dynamics, such as endurance and speed, are affected by neonicotinoid exposure. However, to date only two studies (both using tethered honeybees) have specifically tested this and have reported mixed findings. One study found acute neonicotinoid exposure increased flight endurance, with the opposite effect shown following chronic exposure (Tosi et al., 2017). The other study detected no effect of chronic exposure on flight performance unless it was provided to individuals in combination with the parasitic varroa mite (Blanken et al., 2015). Hence, further investigation is needed to understand the generality of the effects of exposure on bee flight, whilst also: i) ensuring that a concentration within the field realistic range is used; ii) gaining a more in-depth analysis of the dynamics of flight during testing; iii) investigating a representative species of wild bee, given there can be differential responses to pesticide exposure between insect pollinator species (Cresswell et al., 2012; Heard et al., 2017; Rundlöf et al., 2015); and iv) considering variation in worker body size, given this can modulate flight capability, can be associated with variation in foraging behaviours within bumblebee colonies (Goulson et al., 2002; Spaethe and Weidenmuller, 2002), and size-specific energetic demands show a non-linear relationship (Greenleaf et al., 2007; Kaufmann et al., 2013).

We investigated the effect of acute oral neonicotinoid exposure on different aspects of bumblebee (*Bombus terrestris audax*) flight performance using a controlled tethered flight mill setup. For this study we exposed individual workers to the neonicotinoid imidacloprid at a concentration of 10ppb as it is: i) a widely used insecticide across the globe with a growing market in many regions (Casida, 2018; Cressey, 2017; Domenica et al., 2017; Mitchell et al., 2017; Zhang, 2018); ii) a concentration that can be found inside social bee colonies, on return foraging workers, and in the pollen and nectar of individual flowers (Blacquière et al., 2012; Botías et al., 2016; Cresswell, 2011; Dively and Kamel, 2012; Goulson, 2013; Hladik et al., 2016); iii) known to impair foraging performance after exposure (Godfray et al., 2014; Pisa et al., 2017); and iv) a neonicotinoid under current scrutiny by policy makers and regulators (Cressey, 2017), resulting in a recent EU ban from agricultural use outside of closed greenhouses. Here, we tested the propensity of individual bees to fly, followed by measures of their flight distance and duration, the dynamics of velocity over the course of the flight test, and investigated how neonicotinoid exposure interacted with worker body size on these performance measures.

## Methods

### Bee husbandry

Three bumblebee *Bombus terrestris audax* colonies, containing a queen and between 130-150 workers, were supplied by a commercial company (Agralan Ltd). Each colony was delivered in a separately housed plastic nest box (29 x 22.5 x 13 cm) and kept in a controlled environment room (25°C) under red light. From the point of arrival, colonies were provisioned with 4g of pollen daily and supplied with ad libitum 10/90% sucrose/water solution via a connected reservoir. A 10% sucrose concentration falls within the range of many flower species (Pierre et al., 1999; Pyke and Waser, 1981), but our primary justification for using this concentration was to ensure individual workers were motivated to feed to satiation when provided a higher concentration of sucrose solution during the acute exposure setup, whilst ensuring that provisioned colonies did not suffer from dehydration or starvation.

### Flight Mill Setup and Bee Tethering

Six flight mills were set up in a separate adjoining room under the same environmental conditions as the housing room (constant 25°C temperature), but with the option to switch between red (Philips TLD 58W Red 1SL/25; mean 660 λ nm) and white light (Philips TLD 58W 840). Flight mills were adapted from a previous design (Smith and Jones, 2012), consisting of a revolving brass wire, with a magnet hanging from one end designed to attach to a metal tag glued to the bee’s thorax through magnetic attraction (Fig. 1). The revolving brass wire was suspended over a central Delrin rod by the repulsive forces of two magnets, preventing friction during arm rotation to allow fluid motion. The Delrin rod was positioned vertically (90° perpendicular) on a horizontally flat triangular Perspex base. A digital Hall-effect sensor placed on one side of the mill detected each complete revolution by the passage of a neodymium magnet (Fig. 1A) and sent an impulse to a Raspberry Pi computer (model B) via a copper wire connector. From here a Python script recorded the time (in seconds) between each impulse, with each revolution defined as a ‘circuit’ from hereon.

**Figure 1.**
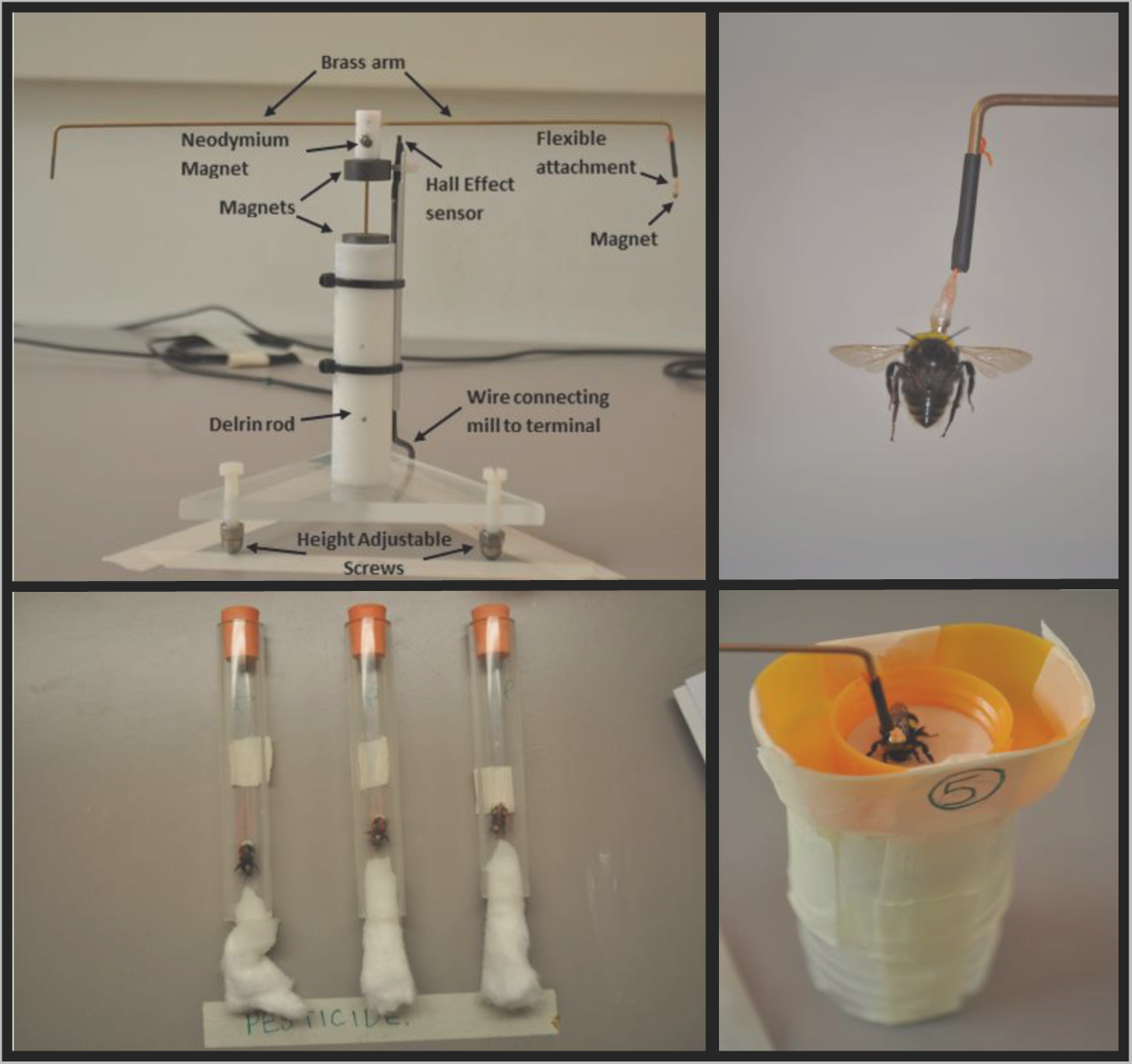
Flight mill setup and associated experimental procedures. Panels show A) flight mill used in the study (the ‘height adjustable screws’ ensured the mill could be horizontal with an attachable bubble level used to ensure this); B) tethering of an individual worker bumblebee to the flight mill magnet; C) feeding procedure in which workers were placed in bunged tubes with one end consisting of cotton wool lightly soaked in 50% sucrose solution (± 10ppb imidacloprid); D) support stands used to hold workers prior to flight tests and following a stop in flight.

When the colonies arrived, we randomly selected 110 workers per colony (total=330) and under red light attached a circular galvanised iron tag (diameter = 2 mm, thickness = 0.4 mm) to the thorax of each worker using super glue (Fig. 2), allowing each individual to be tethered to the hanging flight mill magnet (Fig. 1B,D). We were confident that tag mass would not cause any significant impairment to bee flight performance, as mean (±s.e.m) tag mass was 18 ± 0.3 mg (calculated from weighing 30 tags), equating to just 7.5% of the mean worker wet mass of all individuals tested in this study (240 ± 5 mg). Indeed, bumblebees are capable of carrying > 50% of their own body mass in nectar alone when foraging (Brian, 1954). Each tag was placed at the centre of the thorax, with the tag leading edge touching the back of the first thoracic stripe (Fig. 2). This placement ensured no impediment of wing movement when attached to the flight mill. The meticulous nature of tagging each bumblebee meant that we scored tag positions as 1 = ideal, 2 = unideal or 3 = poor (Fig. 2), with scores 1 and 2 being considered acceptable to experimentally test but score 3 being excluded from further use. In total, 74 workers per colony were tested (total = 222).

**Figure 2.**
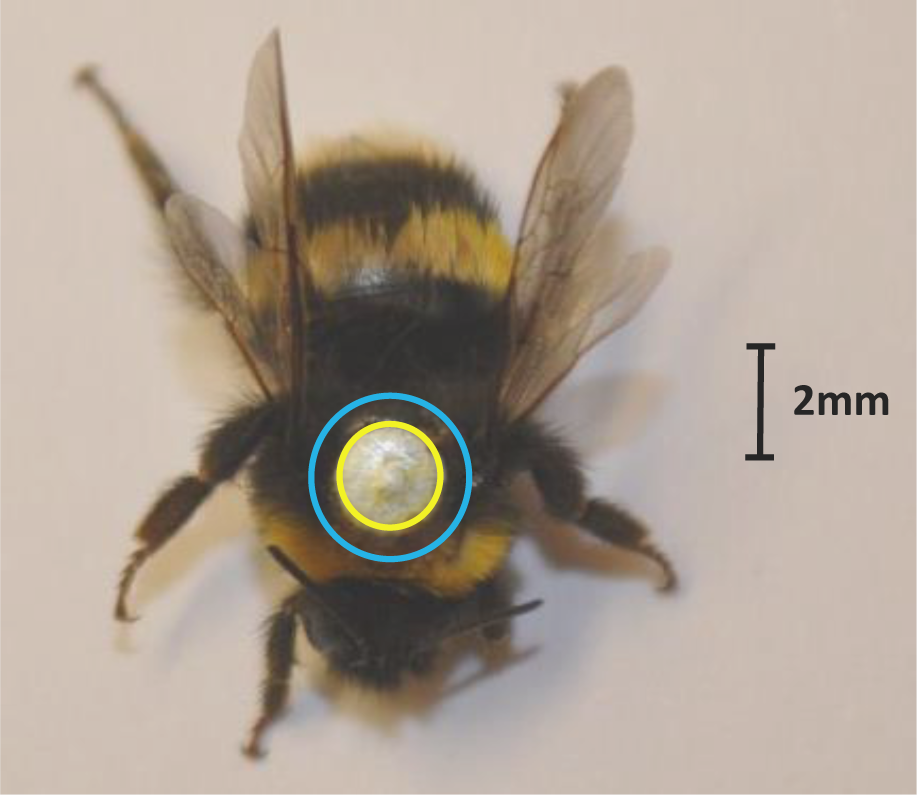
Example of the ideal positioning of a metal tag (tag score 1) on the thorax of a *Bombus terrestris audax* bumblebee worker. If the tag positioning was unideal (tag score 2) the metal tag would overlap the yellow circle but remain inside the blue circle. If positioning was unacceptable (tag score 3) it would overlap the blue circle.

**Figure 3.**
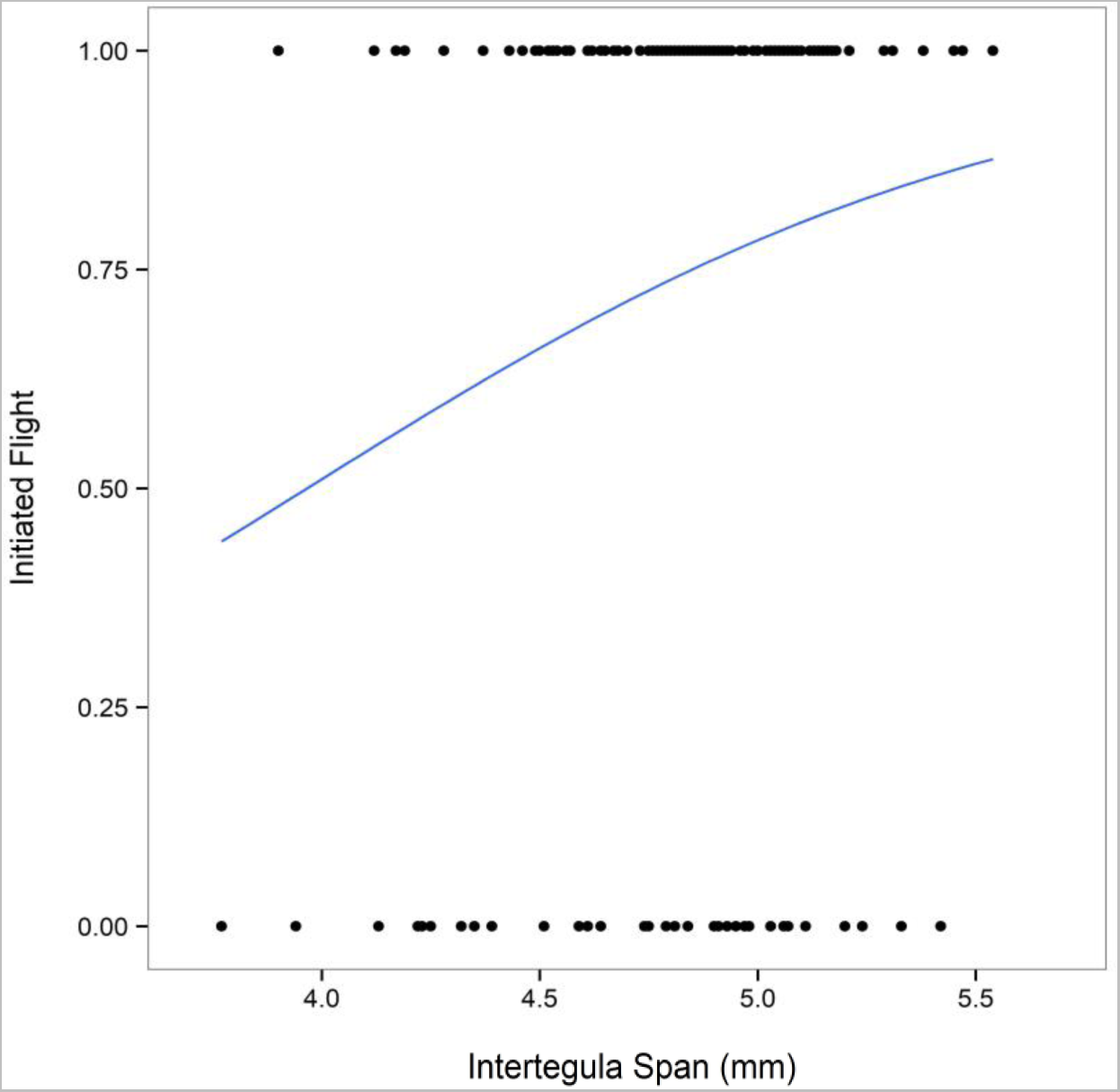
Logistic regression plot showing the effect of body size (*ITS*) on the propensity to fly. All workers from both treatments were pooled (n=140) and could either have initiated flight (= 1) or refused to fly (= 0; Table 1 – filter step 4).

**Table 1.**
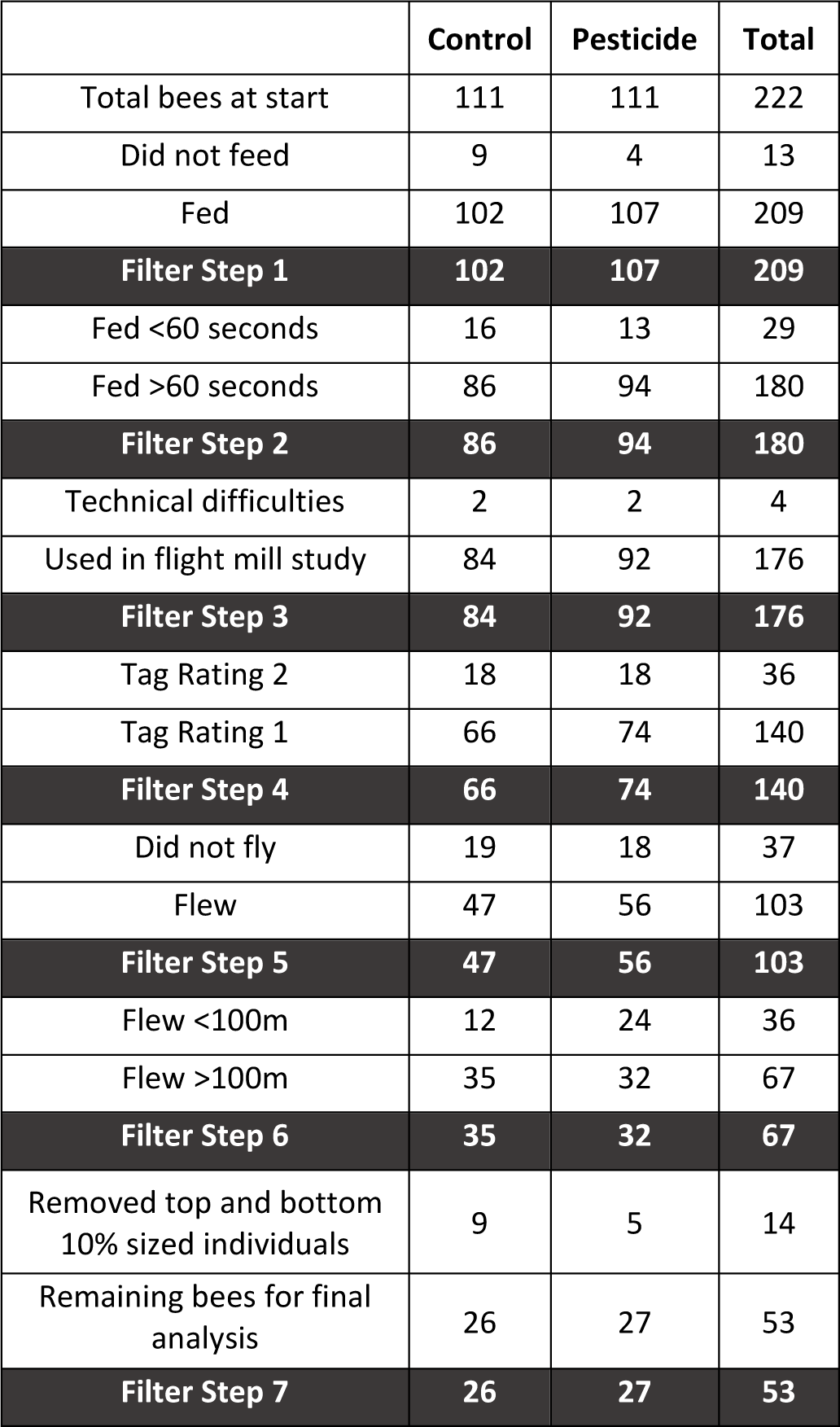
An overview of the filter steps used when cleaning the data for analysis of flight performance, outlining the number of workers removed from each treatment at each stage.

### Pesticide Preparation

A 10ppb imidacloprid working solution was produced to supply the acute pesticide exposure to worker individuals. A previously made stock solution of 1ppt imidacloprid dissolved in acetone was used which was stored in a freezer wrapped in aluminium foil to prevent light degradation (Soliman, 2012). Aliquots of the stock solution were diluted by addition of a 50% sucrose solution to create the working 10ppb *pesticide* treatment solution. The control solution was made by adding the respective volume of acetone to the 50% sucrose solution to produce a 10ppb acetone *control* working solution.

### Experimental Procedure

Pilot studies were first conducted in September 2015 and March 2017 to trial and verify the experimental setup and procedures, with the main experimental study conducted in April 2017. The main experimental testing started 12 hours after tagging was completed, and testing took place over an 8-day period. Workers were tested in bouts, with 5-6 bouts undertaken per day. Six workers were sampled per bout (one per flight mill), consisting of two workers sampled per colony, with one worker randomly assigned to the *treatment* and the other to the *control.* This ensured that all three colonies and both treatment groups were represented equally in each bout and over the totality of the experiment (total: n = 37 bees per treatment per colony; n = 111 *control* & 111 *pesticide*). Once removed from the colony, each worker was directly transferred to a separate transparent horizontally laid Perspex tube (length = 150mm, ID = 19mm). The tube had a rubber bung at each end creating a holding compartment for the bee, with individuals left to acclimatise for a resting period of 3 minutes. After this resting period, one of the bungs was replaced with cotton wool lightly soaked in the *control* or *pesticide* treatment sucrose solution (Fig. 1C). This already piloted method ensured that 94.1% of workers in our main experiment fed. We made the assumption that spiking the sucrose solution with the neonicotinoid would not deter feeding if given no other option, as supported by pilot observations and previous studies (Arce et al., 2017; Gill et al., 2012; Kessler et al., 2015). Our pilot study also indicated that workers took a mean (±s.e.m) duration of 50 ± 13 seconds to commence feeding (defined as prolonged proboscis extension on to the cotton wool) and fed for a mean (±s.e.m) duration of 213 ± 24 seconds before stopping, with subsequent feeds being rare, sporadic, and short (<10 seconds). Workers could access the provisioned sucrose-soaked cotton wool for 10 minutes, after which the cotton wool was removed and original bung replaced, followed by a 5-minute resting period inside the tube. Whilst this protocol meant that we could not determine the precise dosage of imidacloprid consumed by each worker, it did allow workers to feed to satiation which is a state likely to occur in the field during foraging bouts, and importantly allows consumption volume to vary proportionately to individual worker size (Free and Butler, 1959; Goulson et al., 2002)

The workers that fed were then removed carefully using tweezers and tethered to the flight mill. The 5.9% of workers that did not feed were immediately frozen (−20°C) and weighed along with all other bees after all flight tests had been completed. All of this was carried out under red light conditions, but once workers were tethered to the mills the room was switched to white light. Each mill had a separate height-adjustable stand which was erected once the bee was tethered and used to hold the worker in place (Fig. 1D). Prior to initiating the flight test, workers were held in place for a period of 10 minutes for two primary reasons: i) pilot observations demonstrated that some bees were initially irritated by attachment to the mill which discouraged flight, but that a 10 min acclimatisation period allowed irritation to subside; ii) a balance was sought between giving workers time to metabolise the neonicotinoid and preventing de-motivation to fly by having them separated from their natal colony for too long. Studies have shown that honeybees metabolise imidacloprid and other neonicotinoids quickly, with a 100µg kg^-1^ dose of imidacloprid showing the greatest levels of presence in the thorax and abdomen after just 20 minutes from ingestion (Suchail et al., 2004), and >50% of a 100µg kg^-1^ dose of acetamiprid being metabolised in less than 30 minutes (Brunet et al., 2005). In our study, a total of 25 mins passed from starting the feeding trial to starting the flight test, which we are therefore confident represents enough time for absorption and metabolism of some of the imidacloprid consumed.

Immediately after the 10-minute acclimatisation period, the support stand was removed quickly from beneath the bee in order to stimulate flight. Prior to removal, the stand was rotated to ensure the worker had a forward-facing orientation. Stand removal caused loss of tarsal contact with the stand surface, which can trigger flight as evidenced in our pilot and other previous studies (Blanken et al., 2015; Brodschneider et al., 2009; Tosi et al., 2017). However, if the worker did not initially start flying the flat side of the stand was used to gently tap the legs in order to generate a sharp loss of tarsal contact. Up to three taps were allowed in this first flight attempt, with individuals being removed from the flight test if no flight was initiated.

Workers that successfully flew in the first attempt were monitored for any subsequent flight stoppages. Each stop was noted down as ‘a strike’, and each worker was permitted five strikes before their flight test was terminated. Immediately following a strike, the individual would be held in the stand to ensure tarsal contact for a 20 second rest period before removal of the stand again. Therefore, in the subsequent data analysis, flight stoppages were identified as circuits with a duration >20 seconds. After a strike, workers were only permitted one tap of the legs in attempt to trigger flight, otherwise their flight test was terminated. Any stoppages that occurred on the first circuit were discounted as genuine stoppages, as this was deemed an acclimatisation circuit for bees to familiarise themselves with the experimental setup. All workers were given the opportunity to fly for up to 60 minutes, including all stops, after which the flight test was terminated. Each individual worker was allowed a maximum of five ‘strikes’, essentially allowing each bee five chances to continue its flight until the 60-minute end point. We felt this provided a better representation of field conditions and a more realistic prediction of foraging distances, as foraging bees do not fly continuously during foraging bouts but will stop at flowers periodically to feed and rest (Woodgate et al., 2016). Additionally, it allows individuals to acclimatise to the conditions of the flight mill and decreases the possibility of excluding individuals from testing that are initially demotivated to fly due to the experimental set up.

Following each flight test, workers were placed in separate labelled tubes and frozen (−20°C). After completion of the whole experiment, for each individual worker we measured: i) wet body mass (including the attached metal tag); and ii) intertegula span (ITS) taken using a digital calliper (Workzone 150mm), with the mean of three repeated measurements being used. For our data analysis, ITS was taken as a proxy for worker body size (Cane, 1987; Greenleaf et al., 2007). This is more appropriate than considering worker wet mass, as wet mass will vary according to both the volume of sucrose solution consumed and the duration of flight, as individuals gain mass through feeding and lose it through energy metabolism during flight.

### Data Cleaning

Frequency distribution plots revealed a spike in the number of workers that terminated the flight test before completing 100m (118 circuits; Fig. S1A). Workers that did not fly over this threshold distance were excluded from the endurance and velocity analysis as a precautionary measure to discount individuals whose flight mill performance is not representative of actual flight capacity. For each worker flying beyond the 100m threshold, we calculated the following: i) total distance flown during the flight test, by taking the total number of circuits flown multiplied by the circuit circumference (0.848m); ii) total duration of the flight test, by summing all circuit interval times; and iii) velocity of each circuit, by taking the circuit circumference and dividing it by the respective circuit interval time. We took a simple calculation for mean velocity, calculated as the total distance flown divided by the total duration flown, and maximum velocity was taken from the circuit showing the highest velocity attained across the flight test.

The velocity calculations for each individual flight test were carried out on cleaned data in which the following circuits were excluded from the analysis: i) first five circuits of the first flight attempt; ii) first five circuits directly following a strike; iii) the circuit directly preceding a strike circuit. It was noted from pilot observations and the main study that removal of the support stand or tapping of the legs would often stimulate strikingly high velocities. It is likely this behaviour is a reaction to stimulatory stress, so we felt actions i) and ii) were justified as a precautionary measure to ensure we only considered circuits representative of normal continuous flight. Similarly, in justification of action iii), the minimal rotational resistance of the mill means that when a worker stops flying it does not equate to an abrupt stop, but the brass arm continues to rotate and slows gradually.

### Data and Statistical Analysis

When considering total duration flown it was noted that the data was bimodally distributed (Fig. S1B), therefore we converted the results to a binary response variable categorised as having or having not flown >2000 seconds, with this duration value decided on as it fell at the bottom of the bimodal concave.

Statistical analyses were conducted using the ‘lme4’ (Bates et al., 2015) package in R v3.2.0 (R Core Team, 2015), with summary statistics generated using the package ‘psych’ (Revelle, 2015) and results reported using the package ‘lmerTest’ (Kuznetsova et al., 2015). A linear model was used to compare variation in *ITS* (body size) between treatments, with *treatment* (*control* or *pesticide*) as the only fixed effect. For all other analyses, mixed effects models (fitted by maximum likelihood) were initially used with *colony* included as a random effect. Unless otherwise stated, fixed effects in each analysis included *treatment* (*control* or *pesticide*), *ITS* and the associated interaction term. Where response variables were binary (propensity to feed, propensity to fly, flight over 100m, flight longer than 2000 seconds) the data were analysed using a GLMM function under a binomial family distribution, with an LMM function used for all other responses (feeding time, total distance flown, mean velocity, maximum velocity). However, where the random effect of *colony* was found to explain none of the variance in the data, it was removed from the model to simplify, and the model reverted to either a GLM or LM instead (the type of model used is indicated with each result). To examine whether an unideal (score 2) tag fitting inhibited flight or impeded movement, we compared the propensity to fly (GLM) and distance flown (LM) between tag ratings for both *treatment* groups separately. Here, the fixed effects were *tag rating* (score of 1=ideal or 2=unideal), *ITS* and the interaction between the two. Flight velocity over time (considering flight over the first 900 circuits) was analysed using an LMM function with the random effect structure nesting individual bee ID within circuit to account for individual repeated measures over time, and fixed effects including *treatment*, *ITS*, *circuit*, and the interaction term between *treatment* and *circuit.* The model suffered from high Eigen values and had trouble converging when considering all 900 repeated measures, therefore to enhance model fit and convergence we scaled the *circuit* variable and considered the average velocity of every 50^th^ circuit (i.e. each bee had a mean per circuit velocity for circuits 1 to 50, 51 to 100, 101 to 150 and so on) resulting in 18 repeated measures. In all cases, model residuals were plotted to confirm the data met the parametric assumptions of the tests used. Where appropriate, normality tests were used to reveal distributions of the data, and those which appeared non-normal were suitably transformed, with details of these found in Appendix 1.

## Results

### Feeding behaviour

We found no significant effect of treatment on the propensity to feed (n= 9 *control* & 4 *pesticide* workers did not feed; GLM: z=1.29, p=0.20). In concordance with our pilot observations, we found that any feeds following the first were sporadic and short, suggesting workers fed to relative satiety on their first feed. Therefore, we used the length of first feeding time as a reliable proxy for total feeding time. Of the 209 workers that fed, the mean (±s.e.m) time spent feeding was 138 ± 9.0 seconds (n=102) and 127.2 ± 7.8 seconds (n=107) for *control* and *pesticide* workers respectively, with no significant difference between treatments (LMM: t=-0.77, p=0.44; Fig S2). For the flight mill testing, however, we decided to include only those workers that fed for >60 seconds (*control* = 86, *pesticide* = 94, total = 180; Table 1), because: i) visualisation of the plotted feeding times suggested initial feeds <60 seconds were relative outliers (Fig S2); and ii) we wanted to increase the likelihood that each worker had fed to satiation. We found that whilst body size was not a significant predictor of feeding time (LMM: t=1.37, p=0.172), the propensity to feed increased with increasing body size (GLM: z=2.643, p=0.008).

### Flight behaviour

The flight data from 140 of the 180 bees tested on the flight mill were analysed (Table 1), as four workers were not considered due to flight mill technical difficulties, and 36 not considered because unideal (score 2) tag application appeared to affect aspects of flight performance (please see below for justification).

#### i) Effect of tag fitting

The propensity of workers to fly was not significantly affected by tag rating (GLM: *control*; z=-0.98, p=0.33: *pesticide*; z=-0.04, p=0.97), although it is interesting that a higher percentage of tag rating 1 (ideal fitting) workers flew compared to tag rating 2 (unideal fitting) workers in both the *control* (71% vs 61%) and *pesticide* (76% vs 72%) groups. When considering total distance flown, however, tag rating 2 workers flew a significantly shorter mean distance compared to tag rating 1 bees in the *control* group (640 vs 1436 m respectively; LM: t=-2.189, p=0.033), with a similar, although non-significant, trend observed in the *pesticide* group (191 vs 415 m; LM: t=-1.643, p=0.11). Furthermore, we saw similar patterns in other flight metrics with tag rating 2 bees showing lower total duration flown (*control* = 1114 vs 2132 secs; *pesticide* = 272 vs 553 secs) and slower mean velocity (*control* = 0.562 vs 0.657 m/s; *pesticide* = 0.618 vs 0.744 m/s). It was therefore decided to exclude all 36 tag rated 2 workers (*control* = 18, *pesticide* = 18; Table 1) from our analyses, to avoid any artefact results.

#### ii) Initial flight behaviour

Flight was initiated by 103 workers, comprising 71% of *control* (n=47 of 66) and 76% of *pesticide* workers (n=56 of 74), revealing a similar propensity to fly between treatments (GLM: z=0.50, p=0.62, Table S1). Body size was found to be a significant predictor of propensity to fly, with the likelihood of flying increasing with *ITS* (GLM: z=2.163, p=0.031; Table S1). This translated to an estimated probability of *control* workers initiating flight of 0.49, 0.77 and 0.92 for workers with a 4mm, 5mm, and 6mm ITS respectively, with a similar pattern observed for *pesticide* workers (Fig. 3). *Pesticide* compared to *control* workers demonstrated a significantly higher termination of the flight test within the 100m threshold at a proportion of 0.43 (n=24 of 56) vs 0.26 (n=12 of 47; GLM: z=-2.115, p=0.035; Table S1) respectively. For instance, a *control* worker with 5mm *ITS* had an estimated proportion of 0.81 chance of flying >100m, compared to just 0.62 for a *pesticide* worker of the same *ITS*. We further found that larger *ITS* significantly increased the probability of flying >100m (GLM: z=2.318, p=0.020), with no clear significant difference in this relationship between treatments (GLM: *treatment*ITS*: z=1.86, p=0.06)

#### iii) Flight endurance & velocity

Inspection of the 67 bees that flew >100m showed an uneven *ITS* distribution between treatments, with a significant bias of larger *pesticide* workers (mean *ITS* of 4.83 ± 0.05 mm vs 4.99 ± 0.04 mm for *control* vs *pesticide* workers respectively; LM: t=2.382, p=0.020). We therefore took a conservative approach and ran two separate analyses on: i) the full dataset including all 67 bees (*control* = 35, *pesticide* = 32); and ii) a subset of the data (*control* = 26, *pesticide* = 27) in which we attempted to normalise the worker *ITS* distribution by removing the smallest 10% (n = 6 *control & 1 pesticide*) and largest 10% (n = 3 *control* & 4 *pesticide*) of workers; resulting in no significant difference in worker *ITS* between treatments (4.86 ± 0.03 mm vs 4.94 ± 0.03 mm for *control* vs *pesticide* workers respectively; LM: t=1.77, p=0.08; Table 1). Normalising the dataset allowed us to better meet the assumptions of our implemented linear models, therefore here we present the analysis using the data subset, and provide the results using the full dataset in the supplementary material (Fig. S3 & Table S2), which showed the same directional pattern on flight performance between treatments.

*Pesticide* workers flew a significantly lower mean (±s.e.m) total distance at just 659.1 ± 78.7 m compared to 1,833.9 ± 207.6 m for *control* (LMM: t=-5.618, p<0.001; Fig. 4A, Table S2). The effect of pesticide exposure on distance flown was mirrored in the effect on duration flown, with a mean (±s.e.m) flight duration of just 822.0 ± 90.8 seconds for *pesticide* exposed workers being considerably shorter than 2,852.2 ± 234.4 seconds for *control* workers (Fig. 4B, Table S2). Visualisation of the durations flown across all workers (Fig. 4B) showed a striking difference between treatments, with a proportion of just 0.04 of *pesticide* workers flying >2000 seconds, whilst 0.81 of *control* workers surpassed this duration (GLMM: z=-4.016, p<0.001, Table S2). Furthermore, a proportion of 0.65 of *control* workers flew for the full 60 minutes permitted, whereas critically not one *pesticide* exposed worker achieved this.

**Figure 4.**
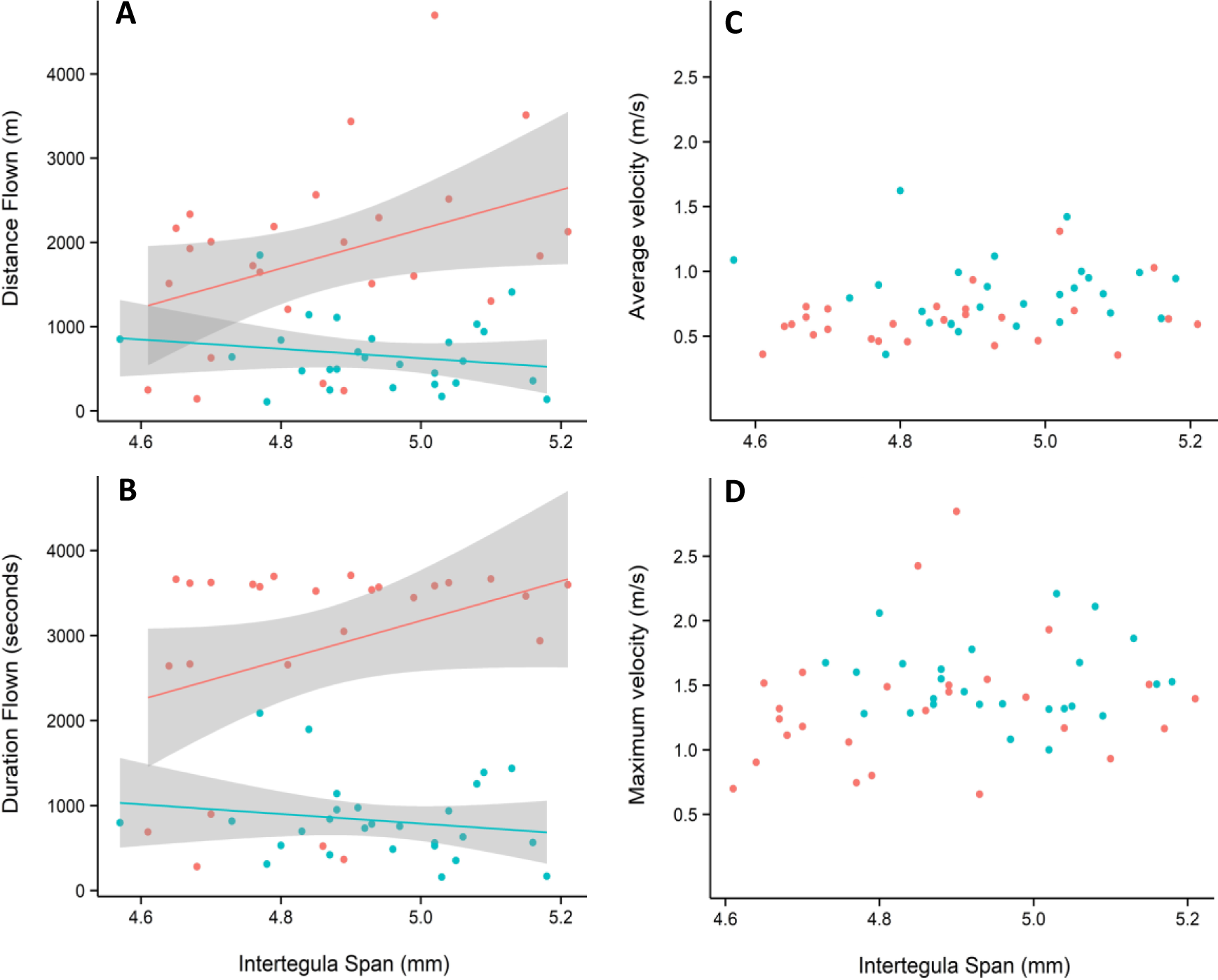
Scatterplot showing key flight performance indicators of endurance (distance flown in meters (A); duration flown in seconds (B)) and average and maximum velocity in meters per second (C-D) against worker body size (*ITS*) for both the *control* (red) and *pesticide* treated (blue) groups. Data plotted is for the subset of bees with normalised *ITS* between treatments (number of workers = 26 *control*; 27 *pesticide*), and linear fitted lines with 95% confidence intervals are estimates of the mixed effects models.

Interestingly, the effect of worker body size on distance flown appeared to differ between *pesticide* and *control* groups, as indicated by a significant *treatment*ITS* interaction (LMM: t=-2.242, p=0.029; Fig. 4A, Table S2). Separate analysis of each treatment group showed that whilst increasing *ITS* resulted in significantly higher total distances for *control* workers (LMM: t=2.158, p=0.041), this relationship was not found for *pesticide* exposed workers (LMM: t=-1.03, p=0.31; Fig. 4A). The effect of *ITS* on total duration flown showed the same general trend as that found for distance (Fig. 4B, Table S2), however, the difference in effect between treatments was less strong (GLMM: z=-1.720, p=0.085). Separate analyses for each treatment group found no significant relationship between *ITS* and the proportion of bees flying >2000 seconds for both treatments (GLMM: *control*: t=1.50, p=0.13; *pesticide*: t=-1.13, p=0.26).

When considering the velocity of individuals across the total flight period, we found *pesticide* exposed workers attained a significantly higher mean (± s.e.m) velocity of 0.84 ± 0.05 m/s per worker compared to 0.63 ± 0.04 m/s for *controls* (LMM: t=2.954, p=0.005; Fig. 4C, Table S2). Looking at maximum velocity, whilst we found no significant difference between treatments (LMM: t=1.58, p=0.12; Fig. 4D, Table S2), it was intriguing that *pesticide* exposed workers were again faster on average (mean ± s.e.m = 1.52 ± 0.06 m/s vs. 1.34 ± 0.09 m/s). This consistent pattern motivated us to examine where these differences in velocity may stem from during flight. Visualisation of velocity over time suggested that *pesticide* workers maintained a higher velocity compared to *controls* during the initial phase (the earlier circuits) of the flight test (see Fig. 5). It also showed a sharp decline in velocity around 900 circuits (760m) as a large proportion of *pesticide* workers terminated flight. Therefore, focusing on the first 900 circuits, we reveal that *pesticide* workers did fly significantly faster compared to *control* workers (LMM: t=3.459, p=0.001; Table S3), with this difference between treatments maintained over these circuits (*treatment*circuit* interaction: t=1.862, p=0.07). Neither mean or maximum velocity was significantly predicted by worker *ITS* (LMM: t=1.60, p=0.12 & t=1.00, p=0.32 respectively; Fig. 4C,D, Table S2), and there appeared to be no effect of *ITS* on velocity over the first 900 circuits (LMM: t=0.50, p=0.62, Table S3).

**Figure 5.**
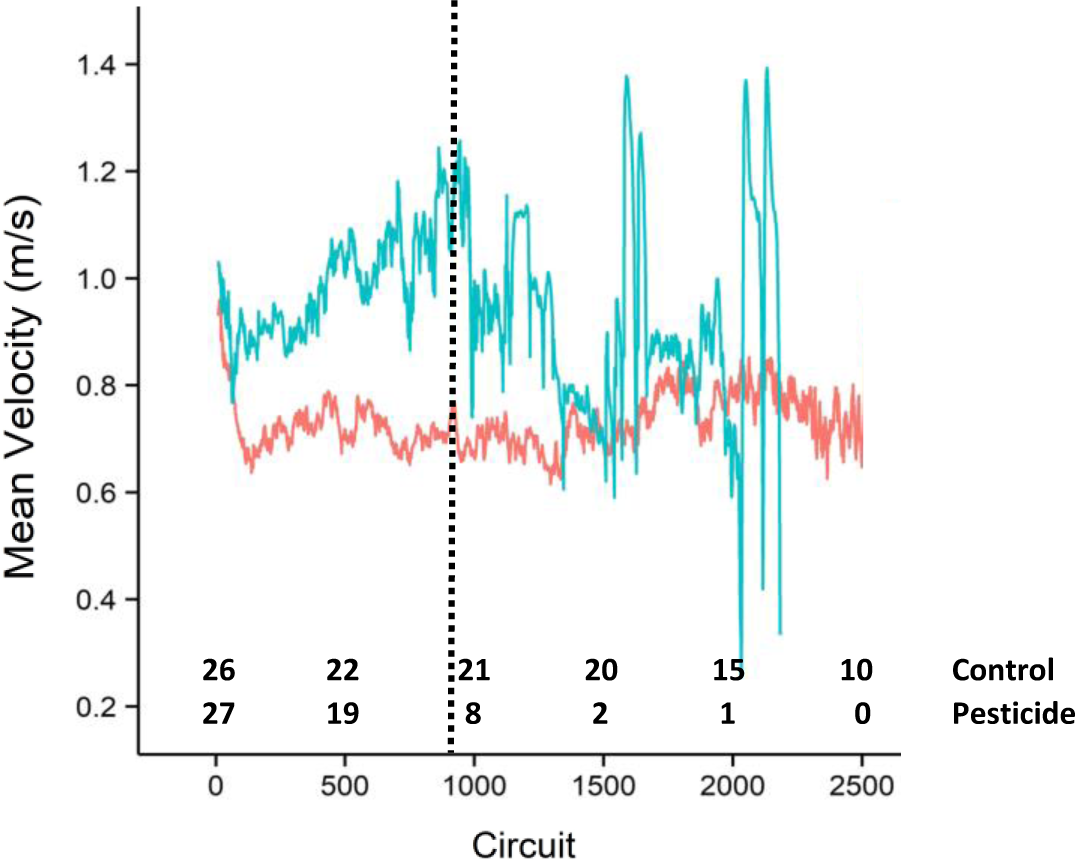
Mean velocity (m/s) flown by each treatment group (*control* = red, *pesticide* = blue) plotted for each consecutive circuit for just the first 2500 circuits. Numbers at the bottom of the graph refer to the number of bees still flying on the corresponding circuit, and the data plotted is for the subset of bees with normalised *ITS* between treatments (starting number of workers = 26 *control*; 27 *pesticide*). Vertical line represents the first 900 circuits used in the analysis for initial individual velocity, and the associated error per mean circuit velocity is not shown.

## Discussion

Despite the importance of bumblebee foraging ability in providing a key pollination service (Garibaldi et al., 2013; Kleczkowski et al., 2017; Stanley et al., 2015), this study, to our knowledge, is the first to test how a specific stressor directly affects the properties of flight in bumblebees. Our findings demonstrate that acute exposure to the neonicotinoid pesticide, imidacloprid, was sufficient to significantly impact overall flight endurance, reducing flight distance and duration to around a third of what *control* workers were able to achieve. Whilst initially both *control* and *pesticide* exposed workers were equally motivated to fly initially, *pesticide* exposed workers showed a higher probability of terminating flight before the end of the 60-minute flight test, which was even evident within the first 100m. Intriguingly, *pesticide* workers exhibited a higher mean velocity compared to *control* workers, which was underpinned by faster flight speeds over the course of the first ¾ km, both during and after which we observed a considerable proportion of pesticide workers terminating their flight. Furthermore, our results suggest that pesticide exposure may negate the capability of larger workers to fly longer distances than their smaller sister workers.

The degree of impact that an acute neonicotinoid exposure had on reducing bumblebee worker flight endurance observed in our study did come as a surprise, as a previous honeybee study showed acute exposure to thiamethoxam increased flight endurance. One possible explanation for these contrasting results is the structural differences between thiamethoxam and imidacloprid compounds, which bind to different sites on nicotinic acetylcholine receptors (nAChRs) with variable affinity (Iwasa et al., 2004; Kayser et al., 2004; Marletto et al., 2003; Wiesner and Kayser, 2000). Indeed, studies have previously shown bumblebees to be less sensitive to thiamethoxam compared to imidacloprid when considering effects on brood production and food consumption (Heard et al., 2017; Laycock et al., 2014). That said, the only other flight mill study to test neonicotinoid effects on honeybee flight capacity also used imidacloprid (as in our study), yet found no effect on workers free from infection with the *Varroa* mite (Blanken et al., 2015). Therefore, this reinforces the view that responses to pesticide exposure can vary considerably even between closely related genera. Indeed, both lab (Cresswell et al., 2012) and field (Rundlöf et al., 2015) studies have highlighted differences in neonicotinoid effects between honeybees and bumblebees, with large interspecific differences in the toxicity of pesticides over time (Heard et al., 2017). Together this emphasises the growing appreciation that reported effects on honeybees cannot always be extrapolated to other wild bees, and highlights the danger of using honeybees as lone indicator species for insect pollinator responses to pesticides (Gill et al., 2016; Heard et al., 2017; Raine and Gill, 2015).

Our flight tests suggest that imidacloprid exposed bumblebee workers experienced a rapid de-motivation to fly as the test progressed and/or tired quickly leading to premature physical exhaustion. Our study was not designed specifically to test these two non-mutually exclusive explanations, however given that only 4% of *pesticide* exposed workers flew >2000 seconds (*control* = 81%) and that not one individual completed the 60-minute test (*control* = 65%), our findings suggest that physical ability may have been affected, which could then have subsequently led to demotivation. We found no difference in initial motivation to fly and in fact *pesticide* exposed workers flew faster than *control* workers, inferring that immediate motor function was not impaired *per se*, but instead flight stamina was reduced. Neonicotinoids have been implicated in affecting honeybee energy metabolism (Derecka et al., 2013), and imidacloprid has been shown to reduce mitochondrial activity, impairing respiratory processes and causing rapid mitochondrial depolarization in neurons of bumblebees and honeybees (Moffat et al., 2015; Nicodemo et al., 2014). Given the high energy expenditure required during flight, a reduction in mitochondrial functioning and the consequent inhibition of ATP production in flight muscles could lead to rapid muscle exhaustion, which might explain our findings of significantly reduced endurance. Imidacloprid can also induce the down-regulation of genes involved in sugar metabolising pathways in honeybee larvae (Derecka et al., 2013), which if true for bee adults could seriously impact flight performance that requires muscles to function at high glycolytic rates (Staples and Suarez, 1997). It is interesting to note that buzz pollination by bumblebees, whereby the creation of resonant vibrations from the flight muscles dislodges pollen from anthers (Morgan et al., 2016), has also been reported to be impaired by neonicotinoid exposure (Whitehorn et al., 2017).

Neonicotinoids are agonists of insect nicotinic acetylcholine receptors (Déglise et al., 2002) and can acutely increase neuronal activity (Matsuda et al., 2001; Moffat et al., 2016). A resultant effect of this may be individual hyperactivity of specific tasks, which could explain our observations of higher velocity in exposed workers during the initial phase of the flight test, and has been previously suggested to underpin neonicotinoid effects on honeybee flight and locomotor activity (Lambin et al., 2001; Suchail et al., 2001; Tosi et al., 2017). Bumblebee colony level exposure to imidacloprid has also been shown to lead to a higher number of workers going out to forage (Gill et al., 2012), a pattern that could be an adaptive response to filling a foraging deficit, but could also be down to possible maladaptive hyperactive behaviour. Additionally, our study suggests a potential cost to hyperactivity, as exposed workers terminated flight prematurely which may have been due to increased energy expenditure during the initial phase leading to faster muscle fatigue and energy depletion, but further testing would be needed to understand this. In sum, our results highlight the importance of looking at the pattern of flight dynamics, rather than experimental end-points, to better understand the mechanisms behind how neonicotinoids act and their temporal effects (Suchail et al., 2001; Wen and Scott, 1997).

Bumblebees have been reported to exhibit a certain degree of alloethism, whereby worker body size can determine divisions in colony tasks (Goulson et al., 2002; Herrmann et al., 2018; Peat et al., 2005). Larger workers of a colony are considered more likely to become committed foragers (Jandt and Dornhaus, 2009; Spaethe and Weidenmuller, 2002), and there have been reports of foraging rate, distance and efficiency (nectar collected per unit time) increasing with body size (Goulson et al., 2002; Greenleaf et al., 2007; Jandt and Dornhaus, 2009; Kapustjanskij et al., 2007; Spaethe and Weidenmuller, 2002; Worden et al., 2005). Whilst our study found no clear relationship with flight velocity and body size, we did find that both the propensity to fly and total flight distance were positively related in *control* workers, which might provide a mechanistic explanation as to why foragers tend to be the larger colony workers. Critically, however, we found no such significant relationships in *pesticide* exposed workers, suggesting that the negative effect of neonicotinoid exposure on flight actually increased in magnitude as workers increased in body size. Intriguingly, a previous study showed that neonicotinoid induced impairment to spatial learning behaviour in bumblebees appeared to be exhibited more highly in the largest colony workers (Samuelson et al., 2016). Together these findings raise the question as to whether larger bumblebees are more susceptible to pesticide effects. With pesticide exposure seemingly counteracting the increased flight performance with body size, the production of larger bees could be seen as wasted energetic investment for the colony. Further investigation is required to look at this, however, as whilst the interactive effect of pesticide and body size was detected in the subset of workers analysed, this effect seemed to be lost when considering the full dataset; a discrepancy that may stem from biases in worker size between treatments as a consequence of the flight trial filtering process.

Bumblebee foraging ranges are difficult to accurately measure, and further knowledge of this important behaviour is critical for predicting colony success and pollination services in changing landscapes. Our flight mill setup showed *control* workers to fly a mean total distance of 1.8km, which appears to sensibly conform to other estimates of bumblebee foraging ranges. Estimated foraging ranges using different techniques including harmonic radar (Osborne et al., 1999), mark-recapture (Kreyer et al., 2004; Osborne et al., 2008), and use of microsatellite genetic markers (Darvill et al., 2004) for *Bombus terrestris* vary from 0.34 to 2.2km. As bumblebees are central place foragers, foraging trips require not only reaching a resource, but also returning to the nest site after collection of food or other resources. The minimum round-trip flight distances associated with the above foraging ranges would therefore span from around 0.68km to 4.4km. Given that our measures fall in the middle of these estimates, we are confident that our flight mill test setup can provide us with meaningful insights into the effects of stress on flight capabilities that can occur in the field. With imidacloprid exposure reducing total flight distance by nearly 1.2km on average, this corresponds to a 64% reduction in comparison to the *control,* which would lead to a notable 87% decline in the total foraging area accessible to a colony (using the colony as the epicentre). Pesticide exposure will therefore place increased stress on bumblebee colonies, with foragers potentially being unable to reach resources they previously could, or unable to return to the nest following exposure feeding on contaminated flowers. Not only would this reduce the abundance, diversity, and nutritional quality of food available to a colony, but could also reduce the pollination service the colony is able to provide (Blanken et al., 2015; Tosi et al., 2017; Van der Sluijs et al., 2013). Looking at the effects of chronic exposure would provide further insights, as bees in the wild would likely be exposed to treated or contaminated flowering plants throughout the season (Simon-Delso et al., 2015; Stanley et al., 2013; Tison et al., 2016).

## Contributions

RJG conceived the project; DK analysed the data; HC, IP, ARR & SG developed the experimental setup; HC performed the experiment; DK, HC and RJG wrote the manuscript.

## Acknowledgements

We are grateful for the advice of Andres Arce, and the valuable assistance of Paul Beasley, Dennis Wildman, Jim Culverhouse and Dylan Smith.

## Experimental Animals

All procedures involving experimental animals were performed in compliance with local animal welfare laws, guidelines and policies.

## Appendix 1

### Statistical Analysis

In all cases, model residuals were plotted to confirm the data met the parametric assumptions of the tests used, and model fit was assessed and optimised where possible. Where appropriate, normality tests were used to reveal distributions of the data, and those which appeared non-normal were suitably transformed as outlined below:

#### Feeding time

The feeding time response variable was box-cox transformed (Venables and Ripley, 2002) to the optimal exponent.

#### Effect of tag fitting on flight behaviour

The effect of tag fitting on flight behaviour was examined separately for both treatments. The effect of tag fitting on total distance flown was analysed using a linear model, with the response variable being square root transformed when considering the *pesticide* treatment, but being left untransformed when considering the *control* treatment.

#### Average and maximum velocity

Prior to maximum velocity analysis, one outlier was identified in the *pesticide* treatment group with a maximum velocity of 14 m/s. This value was around 12 m/s higher than any other individual, and being considered a reaction to stimulation as opposed to normal flight behaviour this value was removed from further analysis. When considering the full dataset of 67 bees (Table 1 – filter step 6), the average velocity and maximum velocity values were square root and cube root transformed respectively. When only considering the subset of 53 bees (Table 1 – filter step 7, both the average velocity and maximum velocity values had to be square root transformed.

#### Total distance flown

For both the full dataset and the subset analysis, the total flight distance response variable was square root transformed.

